# Metabolic engineering of *Escherichia coli* for optimized biosynthesis of nicotinamide mononucleotide, a noncanonical redox cofactor

**DOI:** 10.1101/2020.05.11.089011

**Authors:** William B. Black, Derek Aspacio, Danielle Bever, Edward King, Linyue Zhang, Han Li

## Abstract

**Background:** Noncanonical redox cofactors are emerging as important tools in cell-free biosynthesis to increase the economic viability, to enable exquisite control, and to expand the range of chemistries accessible. However, these noncanonical redox cofactors need to be biologically synthesized to achieve full integration with renewable biomanufacturing processes.

**Results:** In this work, we engineered *Escherichia coli* cells to biosynthesize the noncanonical cofactor nicotinamide mononucleotide (NMN^+^), which has been efficiently used in cell-free biosynthesis. First, we developed a growth-based screening platform to identify effective NMN^+^ biosynthetic pathways in *E. coli*. Second, we explored various pathway combinations and host gene disruption to achieve an intracellular level of ~1.5 mM NMN^+^, a 130-fold increase over the cell’s basal level, in the best strain, which features a previously uncharacterized nicotinamide phosphoribosyltransferase (NadV) from *Ralstonia solanacearum.* Last, we revealed mechanisms through which NMN^+^ accumulation impacts *E. coli* cell fitness, which sheds light on future work aiming to improve the production of this noncanonical redox cofactor.

**Conclusion:** These results further the understanding of effective production and integration of NMN^+^ into *E. coli*. This may enable the implementation of NMN^+^-directed biocatalysis without the need for exogenous cofactor supply.

## Introduction

In the last decade, cell free biosynthesis has emerged as a prominent tool in the production of renewable chemicals, fuels, and pharmaceuticals [1–3] Cell-free systems, both purified enzyme-based and crude lysate-based, have unique advantages over whole-cell biotransformation systems. For example, environmental conditions can be varied within a wider range to favor product formation [4]; transportation issues across cell membranes are eliminated [5]; toxic compounds can be produced at much higher titers than the cell’s tolerance limit [6]. Because components of the biosynthetic pathways can be readily mix-and-matched in a combinatorial fashion, cell-free biosynthesis has also been used as a high-throughput prototyping tool to inform pathway design in whole-cell biosynthesis [7,8].

Cofactors such as nicotinamide adenine dinucleotide (phosphate) (NAD(P)^+^) are essential reagents in biosynthesis. In cell-free biosynthesis, cofactors are freed from the life-sustaining roles they play *in vivo*. Therefore, true opportunities exist to significantly expand the toolkit of cofactors beyond what is offered by Nature to achieve desirable goals in biocatalysis. For example, cheaper noncanonical cofactors, such as 3-carbamoyl-1-phenethylpyridin-1-ium chloride (P2NA^+^) [9,10], have been used in purified enzyme-based redox catalysis to increase economic viability. Noncanonical cofactors with stronger electron-accepting capability, such as 3-acetylpyridine adenine dinucleotide [11,12], have been used to drive the thermodynamically unfavorable reactions of alcohol oxidation.

We recently developed a cell-free biosynthesis platform surrounding the noncanonical redox cofactor nicotinamide mononucleotide (NMN^+^) [13]. NMN^+^ was enzymatically cycled by pairing an engineered glucose dehydrogenase from *Bacillus subtilis* with a variety of enzymes to reduce activated C=C double bonds, activated C≡C triple bonds, nitro groups, and to supply electrons to a P450. This system demonstrated robust temporal stability over 96 hours and a total turnover number of ~39,000. Because of its smaller size, NMN^+^ has also been shown to provide a faster mass transfer rate in enzymatic biofuel cells [14].

Compared to other noncanonical cofactors which are made through chemical synthesis [15–17], NMN^+^ is particularly suited for fully renewable biomanufacturing processes because it is accessible through biosynthesis [18–20]. This feature is especially desirable in crude lysate-based cell-free biosynthesis and whole-cell biosynthesis, where NMN^+^ produced in the cells does not need to be purified or exogenously supplied, and it can be directly used for downstream biocatalysis. Importantly, since we demonstrated its successful application in *E. coli* whole cells to enable orthogonal electron delivery [13], NMN^+^ can potentially be utilized in crude lysate-based biosynthesis to control the flow of reducing power and mitigate side reactions based on the same principles [13,21].

Although NMN^+^ has been biosynthesized previously in metabolically engineered *E. coli* [13,19], further improving NMN^+^ production requires more efficient pathways and a better understanding of its metabolism in the host. While previous efforts have primarily used the nicotinamide phosphoribosyltransferases, NadV, to convert nicotinamide to NMN^+^, only a few NadV homologs have been tested and many other NadV-independent pathways for NMN^+^ biosynthesis remain unexplored. Furthermore, whether and how NMN^+^ accumulation impacts cell physiology remains largely unknown. In this work, we developed a growth-based screening platform to identify pathways for efficient NMN^+^ generation *in vivo*. This platform was designed by making NMN^+^ an essential precursor in NAD^+^ biosynthesis in engineered *E. coli*. We used this platform to demonstrate that NMN^+^ synthetase, NadE* from *Francisella tularensis*, effectively mediates an additional route for NMN^+^ biosynthesis in *E. coli*. We also bioprospected for NadV homologs based on comparative genomic data [18], and we tested their ability to produce NMN^+^ in combination with *F. tularensis* NadE*. The best NMN^+^ producing strain accumulated ~1.5 mM of intracellular NMN^+^ while overexpressing *F. tularensis* NadE* and *Ralstonia solanacearum* NadV simultaneously, as well as harboring a disruption in the gene encoding NMN^+^ amidohydrolase, PncC, Although our current highest NMN^+^ production titer did not cause growth inhibition, we observed inhibitory effect when very high concentrations of NMN^+^ was fed to the cells with an overexpressed heterologous NMN^+^ transporter. Interestingly, we showed that this inhibitory effect can be alleviated when the transcriptional regulator of NAD^+^ biosynthesis, NadR, was disrupted. Together, these results provide insight for future metabolic engineering efforts aiming to further improve NMN^+^ biosynthesis.

## Results

### Identification of NMN^+^ biosynthetic routes

In *E. coli* cells, NMN^+^ is only present at a nominal level, ~11.5 μM as previously reported [13], as the product of the DNA ligase reaction [18]. On the other hand, NMN^+^ accumulates to higher levels and serves as a main intermediate in NAD^+^ biosynthesis in various other organisms [22,23]. Here, we sought to systematically investigate the effectiveness of these heterologous NMN^+^ biosynthetic routes in *E. coli* (Figure 1).

**Figure 1.**
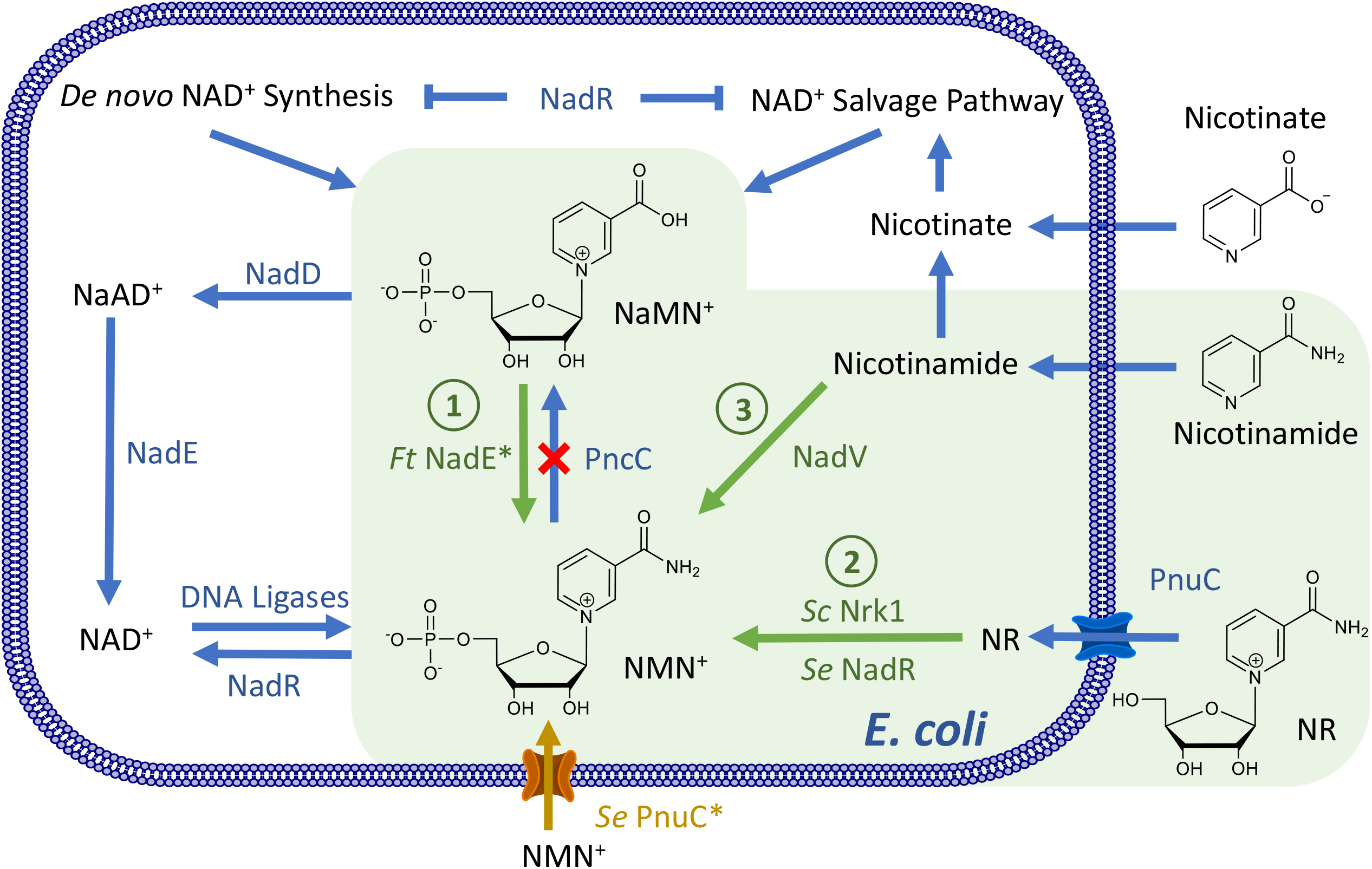
Establishing NMN^+^ Biosynthetic Routes in *Escherichia coli.* NMN^+^ is produced in a small amount through the DNA ligase reaction in the *E. coli* cell. Heterologous enzymes, shown in green, can introduce new routes to generate NMN^+^. *E. coli* endogenous genes and transport processes are shown in blue. Pathway 1 introduces an NMN^+^ synthase from *Francisella tularensis* to produce NMN^+^ from NaMN^+^. Pathway 2, NR salvage, produces NAD^+^ from NR. Pathway 3, NA salvage, produces NMN^+^ from NA. The NMN^+^ transporter PnuC* from *Salmonella enterica*, shown in orange, enabled transport of NMN^+^ into the cell. The endogenous *pncC*, was targeted for gene disruption to prevent NMN^+^ degradation. In the presence of NAD^+^, NadR inhibits transcription of the genes involved in *de novo* NAD^+^ biosynthesis, nicotinate salvage, and *E. coli pnuC*. NaMN^+^, nicotinic acid mononucleotide; NaAD^+^, nicotinic adenine dinucleotide; NAD^+^, nicotinamide adenine dinucleotide; NMN^+^, nicotinamide mononucleotide; NR, nicotinamide riboside; NadD, NaMN adenylyltransferase; NadR, NMN adenylyltransferase; NadE, NAD synthase; *Ft* NadE*, NMN synthase from *Francisella tularensis*; PncC, NMN amidohydrolase; NadV, nicotinamide phosphoribosyltransferase; Pnuc, nicotinamide riboside transporter; Pnuc*, a mutant PnuC enabling direct transport of NMN^+^ across the cell membrane.

Three major NMN^+^ biosynthetic pathways exist in Nature (Figure 1): Pathway 1 produces NMN^+^ from nicotinic acid mononucleotide (NaMN^+^) using NMN^+^ synthetase (NadE*), and it was shown to be part of the *de novo* NAD^+^ biosynthetic pathway in a small group of prokaryotes including *F. tularensis* [23]. Pathway 2 involves phosphorylation of nicotinamide riboside (NR) and functions to salvage NR to ultimately yield NAD^+^. To establish this pathway, we chose to overexpress the native NR transporter in *E. coli*, PnuC [24], in conjunction with two different NR kinases, Nrk1 from *Saccharomyces cerevisiae* [25] and NadR from *Salmonella enterica* [26]. Pathway 3 uses nicotinamide phosphoribosyltransferase (NadV) to convert nicotinamide (NA) to NMN^+^; it plays a role in NA salvage in vertebrates and some bacteria. Marinescu and coworkers demonstrated NMN^+^ accumulation in *E. coli* by heterologously expressing three NadV homologs from *Haemophilus ducreyi*, *Shewanella oneidensis*, and *Mus musculus* while feeding NA [19], and they showed that *H. ducreyi* NadV performed the best [19]. We previously showed that NadV from *F. tularensis*, which belongs to a different clade in the NadV phylogenetic tree to *H. ducreyi* NadV [18], can also effectively produce NMN^+^ in *E. coli* [13]. Here, we sought to explore more bacterial NadVs in the same family of *F. tularensis* NadV and compare them with *H. ducreyi* and *F. tularensis* NadV. Namely, we chose NadV homologs from *R. solanacearum*, *Synechocystis sp.*, and *Synechococcus elongatus* [18].

All three pathways have unique advantages. While Pathway 3 requires a much cheaper substrate than Pathway 2 (NA versus NR), the latter incorporates ATP hydrolysis as a robust driving force. Since NaMN^+^ is an intermediate in *E. coli*’s *de novo* NAD^+^ biosynthesis and can be efficiently produced from central metabolites (Figure 1), Pathway 1 has the potential to achieve complete NMN^+^ biosynthesis from simple feed stocks such as glucose.

### Evaluating the NMN^+^ biosynthetic pathways *in vivo*

We evaluated the three above-mentioned pathways in *E. coli* using a growth-based screening platform (Figure 1, 2). To link NMN^+^ production to cell survival, we employed *E. coli* strain 72c, which contains a temperature sensitive allele of *nadD*, an essential gene in NAD^+^ biosynthesis. As a result, the cells cannot grow at 42 °C [27] unless NMN^+^ can accumulate inside the cells and be converted to NAD^+^ by *E. coli* NadR (Figure 1, 2). Previous work has also established NMN^+^-dependent NAD^+^ biosynthesis to rescue NAD^+^ auxotroph in *E. coli* [28].

**Figure 2.**
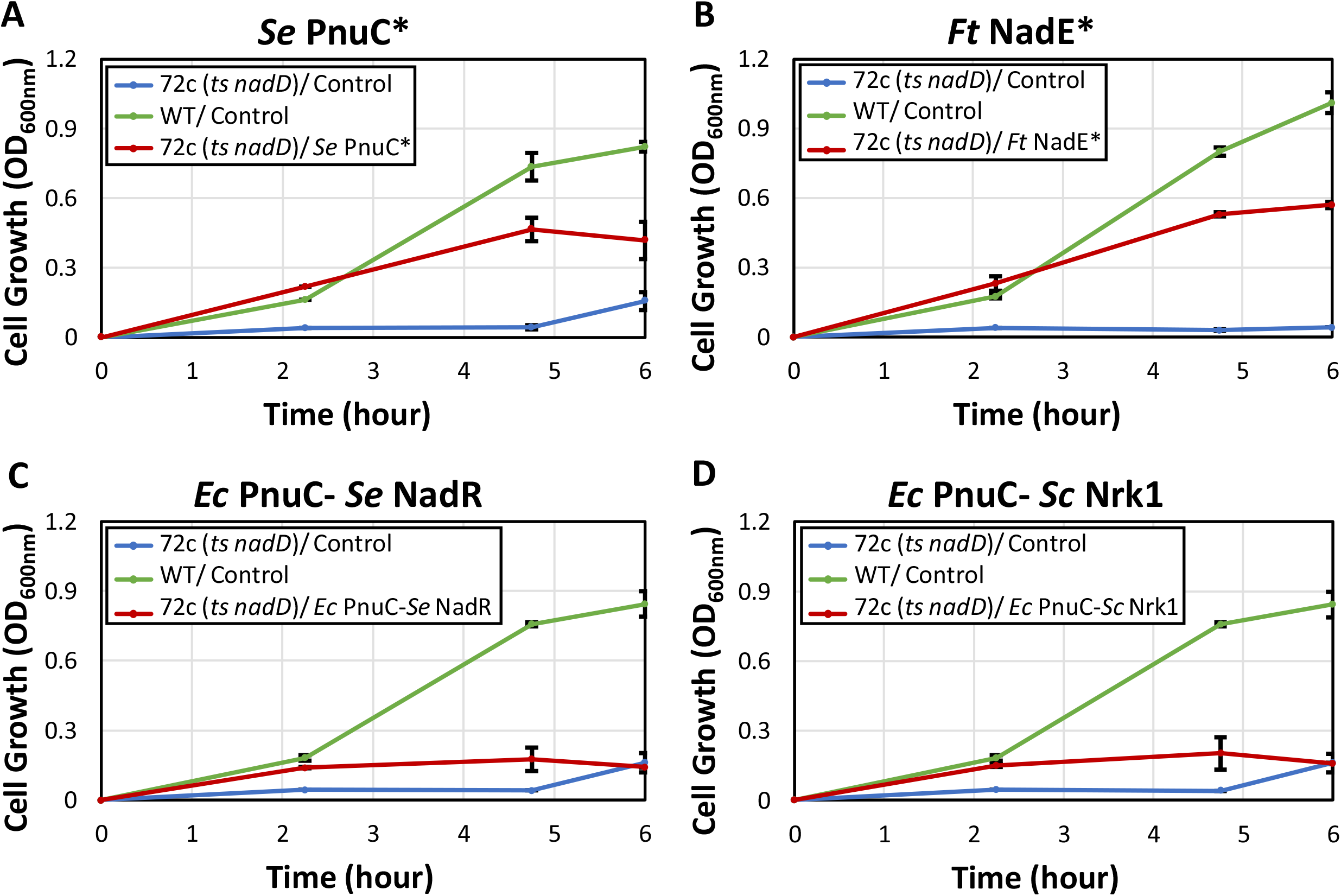
Identification of efficient NMN^+^ production pathways *in vivo* using a growth-based screening platform. A growth-base screening platform was used to identify pathways which efficiently generated NMN^+^ *in vivo*. *E. coli* strain 72c, which contains a temperature sensitive allele of *nadD* (*ts nadD*), which exhibits a conditionally lethal phenotype when cultured at 42 °C because the native NAD^+^ biosynthesis is disrupted. Therefore, the cell must rely on intracellular NMN^+^ to restore NAD^+^ formation and growth. (A) Direct feeding of NMN^+^ into the growth medium with the overexpression of an NMN^+^ transporter, PnuC* from *S. enterica*, restored growth to levels near the wild type control, indicating the platform is effective for NMN^+^ production screening. (B) Introducing *F. tularensis* NadE* also restored growth with the supplementation of nicotinamide. (C, D) Overexpression of *E. coli* ’s native nicotinamide riboside (NR) transporter PnuC, paired with *S. enterica* NadR (C) or *S. cerevisiae* Nrk1 (D) while feeding NR failed to efficiently restore growth. Screening was performed in a deep-well 96-well plate containing 1 mL of LB medium supplemented with 2 g/L D-glucose and 200 μM of NMN^+^ precursors, if applicable.

When 200 μM NMN^+^ was directly fed to the cells expressing an NMN^+^ transporter from *S. enterica* PnuC* [29] (on a multiple-copy plasmid pWB302), growth was restored to levels comparable to wild type cells after 6 hours (Figure 2A). In contrast, cells carrying a control plasmid (pWB301) could not grow, suggesting the basal level of NMN^+^ in *E. coli* cells does not cause background issues in this growth-based screening, possibly because *E. coli* NadR has low affinity towards NMN^+^ [30]. These results demonstrate that the screening platform is functioning properly, and cell growth is serving as a readout for intracellular NMN^+^ level. We and others have previously shown that NMN^+^ can enter *E. coli* cells without overexpressing a heterologous transporter [13, 28]. In this work, the amount of NMN^+^ supplementation can be substantially reduced, suggesting that *S. enterica* PnuC* improves the efficiency of NMN^+^ transportation into *E. coli* cells.

Overexpression of *F. tularensis* NadE* (Figure 1, Pathway 1, on the plasmid pWB303) with 200 μM nicotinamide supplementation restored growth to a similar level as directly feeding NMN^+^ (Figure 2B). Nicotinamide can yield the substrate of NadE*, namely NaMN^+^, through *E. coli* ’s native salvage pathway. Interestingly, growth restoration by *F. tularensis* NadE* does not depend on nicotinamide feeding (Figure S1). NMN^+^ production using S. *cerevisiae* Nrk1 or *S. enterica* NadR (Figure 2, Pathway 2, on the plasmids pWB304 and pWB305) with 200 μM NR supplementation was not efficient enough to restore growth (Figure 2C, D). These results suggest that besides the well-established NadV route (Figure 1, Pathway 3) [19], the *F. tularensis* NadE*-dependent pathway is also effective in *E. coli* for NMN^+^ biosynthesis.

### Bioprospecting NadV homologs and optimizing NMN^+^ biosynthesis

After demonstrating NMN^+^ can be effectively generated by overexpressing *F. tularensis* NadE*, we examined the effects of pairing it with different NadV homologs. *F. tularensis* NadE* was overexpressed in a synthetic operon on a multiple-copy plasmid with each of the five NadV candidates as described above (pWB203, pDB101, pDB102, pDB103, pDB104). Cells were fed 1 mM of NA and grown for four hours before processing and quantification of intracellular NMN^+^ and NAD^+^ levels *via* LC-MS analysis as previously described [13]. When using the wild type BW25113 cells as the host, ~12 to 51 μM of NMN^+^ was produced through these pathways (Figure 3). We previously found that low levels of intracellular NMN^+^ could be attributed to NMN^+^ degradation by the NMN^+^ amidohydrolase, PncC [13]. Expression of the NadE*/NadV pathways in a Δ*pncC* strain, JW2670-1 significantly increased the intracellular NMN^+^ levels. Cells expressing *F. tularensis* NadE* and *R. solanacearum* NadV in Δ*pncC* strain reached the highest intracellular NMN^+^ level of ~1.5 mM, a 130-fold increase over the cell’s basal NMN^+^ level [13], when tested under the same conditions (Figure 3). Furthermore, the *R. solanacearum* NadV strain performed significantly better than *F. tularensis* NadV, the NadV we used in our previous work [13], exhibiting a 2.8-fold increase in intracellular NMN^+^ concentration. *R. solanacearum* NadV also performed better than *H. ducreyi* NadV [19].

**Figure 3.**
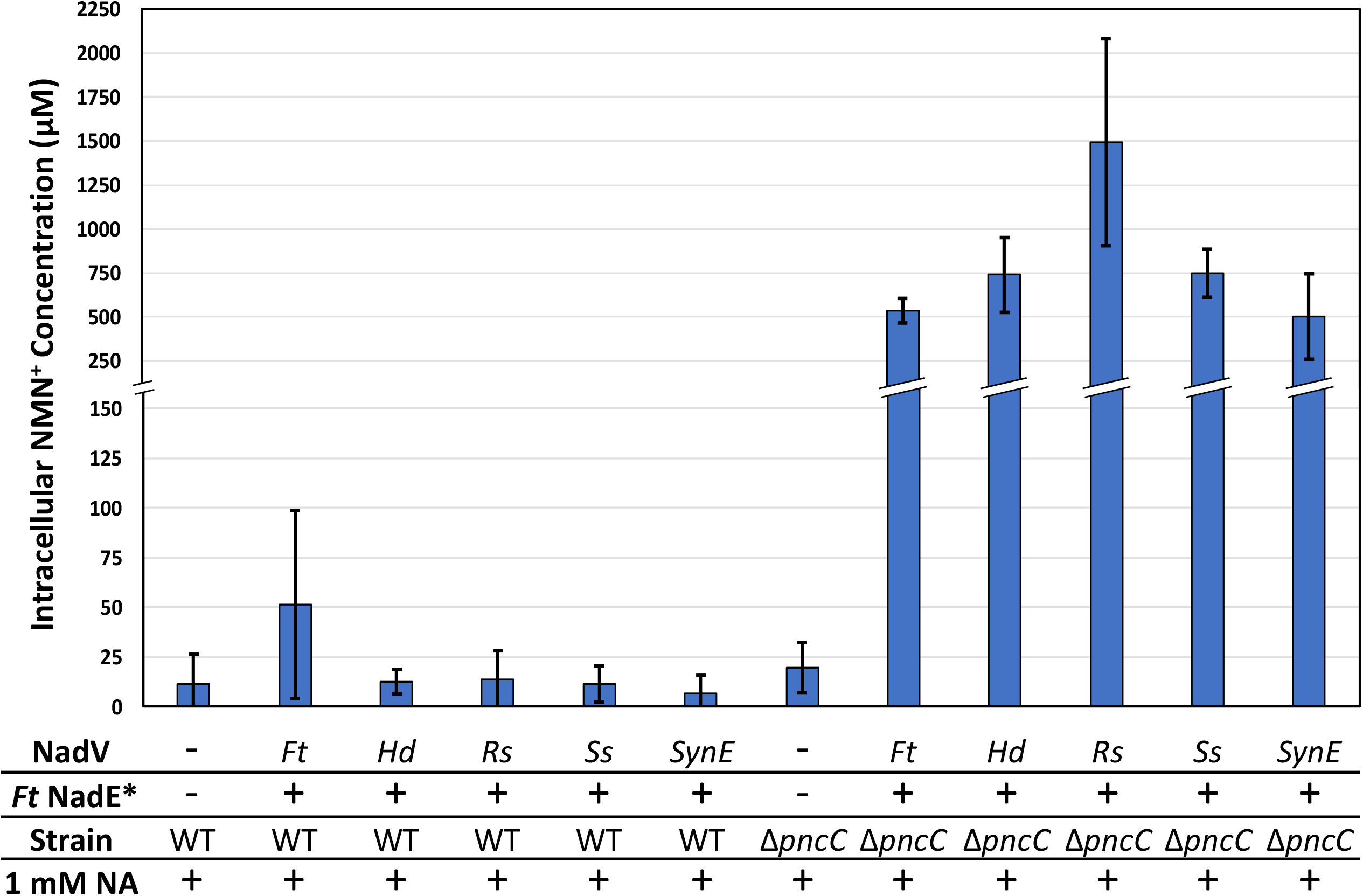
Pathway combination and strain modification improved NMN^+^ production. NadV homologs were co-overexpressed with *F. tularensis* NadE* in wild type and Δ*pncC cells*. In wild type cells, introducing *F. tularensis* NadE* and NadV homologs only resulted in low levels of NMN^+^ accumulation. Disrupting the NMN^+^ degrading-enzyme PncC greatly increased intracellular NMN^+^ levels. Of the NadV homologs tested, *R. solanacearum* NadV demonstrated the highest intracellular NMN^+^ production of ~1.5 mM in Δ*pncC* cells, a 130-fold increase over the cell’s basal level. Cells were grown in 2xYT medium supplemented with 1 mM nicotinamide at 30 °C for 4 hours. NMN^+^ concentration was determined by LC-MS. *Ft, Francisella tularensis*; *Hd*, *Haemophilus ducreyi*; *Rs*, *Ralstonia solanacearum.*; *Ss*, *Synechocystis sp.*; *SynE*, *Synechococcus elongatus*.

### Investigating the physiological response to NMN^+^ accumulation

Even though the *de novo* NAD^+^ biosynthesis pathway was unmodified, the Δ*pncC* cells overexpressing *F. tularensis* NadE* and NadV homologs had lower intracellular NAD^+^ levels compared to the control strain in which no proteins were overexpressed and intracellular NMN^+^ level was low (Figure 3, S2). This suggests that maintaining intracellular NMN^+^ at millimolar-range concentrations may be detrimental to cellular fitness. Although we observed no growth defects in our current best NMN^+^ producing strain, the potential physiological effects of NMN^+^ accumulation may become a bottleneck for future strain optimization.

Since the effects of NMN^+^ accumulation on *E. coli* are not well understood, we sought to stress the cells by drastically increasing the intracellular NMN^+^ levels in the Δ*pncC* strain and observe cellular growth. NMN^+^ concentration was titrated in the medium while overexpressing the NMN^+^ transporter *S. enterica* PnuC*. We found that cell growth was inhibited at high NMN^+^ concentrations (> 5 mM) (Figure 4), which suggests that elevated NMN^+^ level may interfere with physiological processes in *E. coli*. Given the decrease in intracellular NAD^+^ level upon NMN^+^ accumulation (Figure S2), we hypothesized that NMN^+^ may regulate NAD^+^ biosynthesis and we sought to examine whether this regulation was mediated by the transcriptional regulator of NAD^+^ biosynthesis, NadR [31]. Interestingly, when NadR was disrupted, the growth inhibition effect of NMN^+^ was significantly alleviated (Fig. 4). These results suggest that NadR may indeed play a role in the physiological response to NMN^+^ accumulation in *E. coli*. NAD^+^ has been suggested to allosterically modulate NadR’s function [32]. Given NadR’s capability to also recognize NMN^+^ [30], further studies are needed to investigate whether NMN^+^ binding induces conformational change in the DNA-binding domain of NadR and modulates its function as a transcriptional regulator.

**Figure 4.**
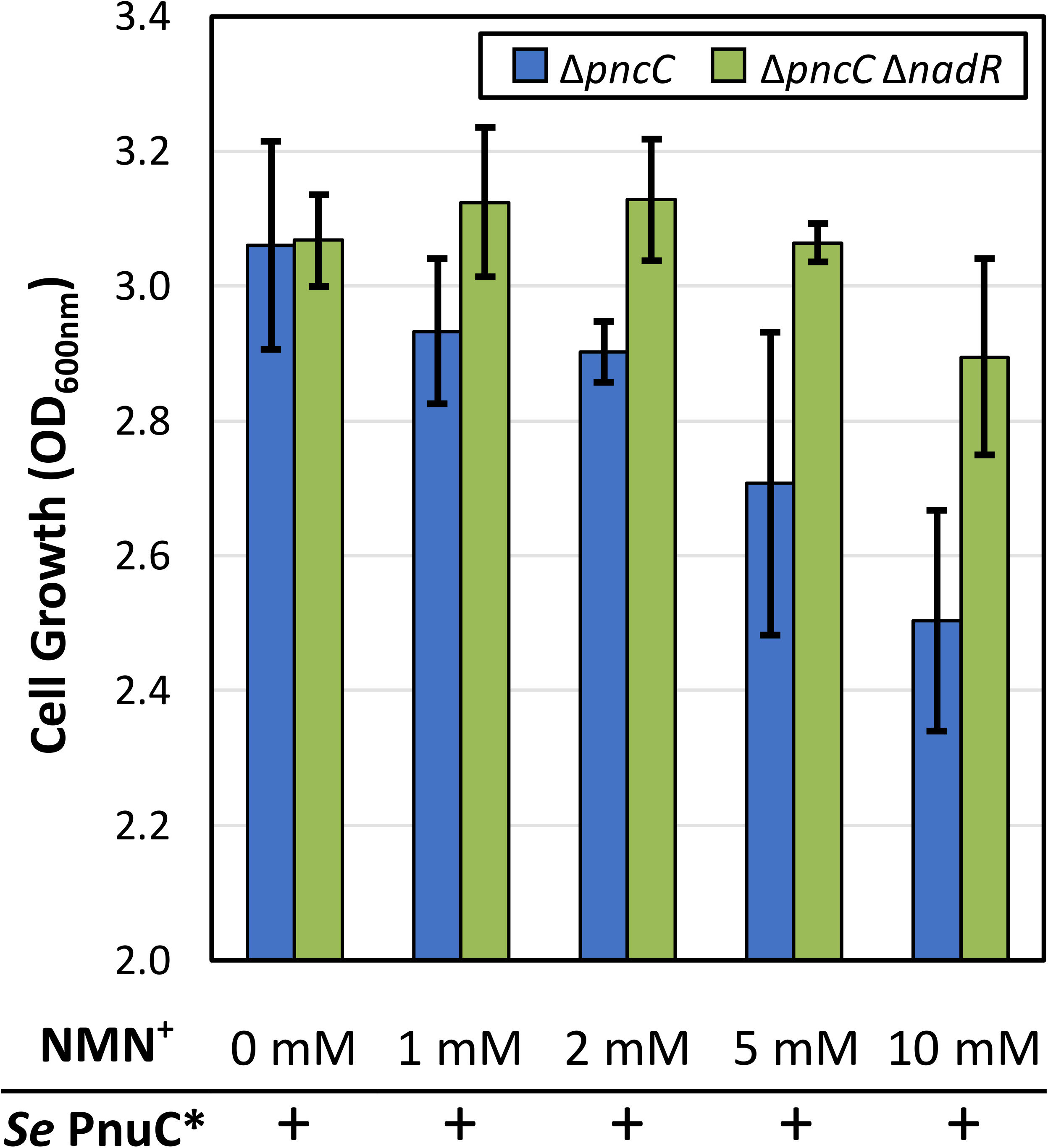
NMN^+^ accumulation affects cell physiology possibly *via* NadR. To determine if NMN^+^ accumulation impacts cell fitness, NMN^+^ was titrated in the growth medium of cells expressing the NMN^+^ transporter *S. enterica* PnuC*. The Δ*pncC* strain exhibited decreased growth at high NMN^+^ concentrations. Disruption of *nadR* significantly alleviated the growth inhibition. This suggests the NadR may mediate the physiological response to NMN^+^ accumulation in *E. coli*. Cells were grown at 30 °C for 6.5 hours in a deep-well 96-well plate containing 1 mL of medium per well.

## Discussion

This work represents the initial steps towards filling some of the fundamental knowledge gaps that remained open in previous work on NMN^+^ biosynthesis in *E. coli*. The important work by Marinescu and coworkers [19] focused on the NadV pathway, but left many other naturally occurring NMN^+^ biosynthetic routes unexplored. Our previous work [13] sought to simply recapitulate the NMN^+^ metabolism of *F. tularensis* [23] by overexpressing both NadE* and NadV from this organism, without dissecting the role of each pathway. Moreover, both efforts did not study the physiological response in *E. coli* to NMN^+^ accumulation. Beside functioning as a cofactor, NAD^+^ is also a universal signaling compound that allosterically controls key enzymes and transcriptional regulators in response to the fluctuating cellular redox state [30,33–36]. Since NMN^+^ is an analog of NAD^+^, it is an open question whether NMN^+^, when it accumulates to a high level, can also interact with the numerous proteins that are modulated by NAD^+^.

The rich information provided by comparative genomic analysis can greatly aid metabolic engineering efforts. By bioprospecting NadV homologs from evolutionarily diverse organisms, we identified *R. solanacearum* NadV, which outperformed the two NadVs that have been previously reported as efficient NMN^+^-producing enzymes in *E. coli* [13,19], when they are compared in the same condition at a bench scale (Figure 3). Since *F. tularensis* NadE* also showed promise to produce NMN^+^ efficiently, similar approach may be taken in the future to bioprospect NadE* homologs.

Moving forward, culture medium and growth conditions can be optimized to potentially yield increased intracellular NMN^+^ levels. In this work, all NMN^+^ production was performed with laboratory standard medium and without optimization. Marinescu and coworkers demonstrated a 32.7-fold increase from 0.72 mM to 23.57 mM of intracellular NMN^+^ upon scale-up to a 500 mL bioreactor while optimizing pH, NA feeding concentration, and dissolved oxygen while culturing in PYA8 medium [19]. Therefore, performing a similar scale-up with our NMN^+^ producing strain may yield significant increases in NMN^+^ production.

In addition, host selection may play a significant role in efficient NMN^+^ biosynthesis. While most industrial model hosts including *E. coli* and *S. cerevisiae* utilize a nicotinic acid adenine dinucleotide (NaAD)-mediated route for *de novo* NAD^+^ biosynthesis, a small group of prokaryotes use NMN^+^ as the primary precursor to NAD^+^ [18,23]. Since NMN^+^ adopts a distinct role and is naturally maintained at a higher level in these organisms [22,23], the physiological responses to intracellular NMN^+^ accumulation may be different. Thus, organisms which utilize NMN^+^-mediated NAD^+^ biosynthesis may be interesting targets for metabolic engineering.

Ultimately, efficient and cost-effective production and purification of NMN^+^ is key for the long-term viability of NMN^+^-based cell-free biotransformation. Once upstream pathways for the renewable production of NMN^+^ are further established, NMN^+^ will need to be extracted and purified before use in cell-free systems. Cells can be isolated through centrifugation, washed, and lysed through homogenization to isolate NMN^+^ from cellular debris. Alternatively, cells can also be permeabilized to release NMN^+^ across the cell membrane, allowing for fewer steps of isolating NMN^+^ from cell mass. Finally, a major advantage of producing NMN^+^ *in vivo* is the direct compatibility with crude lysate-based cell-free and whole-cell biosynthesis. By using cells that are capable of both producing intracellular NMN^+^ and expressing enzymes of interest, crude lysates or whole cells can be directly used for NMN^+^-dependent biosynthesis without the exogenous supply of redox cofactors.

## Conclusions

In this work, we explored routes to efficiently produce NMN^+^ in *E. coli*. After surveying the routes for NMN^+^ production *in vivo*, bioprospecting NadVs enabled the production of 1.5 mM of NMN^+^ using the NadV from *R. solanacearum.* Under the conditions tested, *R. solanacearum* outperformed the previous best NadV’s shown to accumulate NMN^+^ efficiently [13,19]. In addition, this work began to elucidate the physiological effects of NMN^+^ accumulation in *E. coli*. However, further investigation is necessary to maintain productivity as NMN^+^ levels are further increased. Ultimately, advancing noncanonical redox cofactor biosynthesis in microorganisms may enable the application of self-sustained, fully renewable cell-free and whole-cell biocatalysis.

## Methods

### Plasmid and Strain Construction

All molecular cloning was performed in *E. coli* XL1-Blue cells (Stratagene). A summary of strains and plasmids used in this study can be found in Table 1. Plasmids were assembled by Gibson Isothermal DNA Assembly [37]. Polymerase chain reaction (PCR) fragments were generated using PrimeSTAR Max DNA Polymerase (TaKaRa). The method for plasmid construction is described below.

**Table 1:** Strains and Plasmids Used in this Study.

| Strains | Description | Reference |
| --- | --- | --- |
| XL-1 Blue | Cloning strain | Stratagene |
| BW25113 | *E. coli* $\Delta(araD-araB)567$, $\Delta lacZ4787(::rrnB-3)$, $\lambda^-$, *rph-1*, $\Delta(rhaD-rhaB)568$, *hsdR514* | Invitrogen |
| JW2670-1 | BW25113 $\Delta pncC::kan$ | Yale *E. coli* Genetic Stock Center |
| MX101 | BW25113 $\Delta pncC$ $\Delta nadR::kan$ | [13] |
| 72c | *E. coli* F<sup>-</sup>, *lacZ4*, *nadD72*(ts,Fs), $\lambda^-$, *argG75* | [27] |
| Plasmids | Descriptions | Reference |
| pWB203 | *P<sub>LlacOI</sub>::Ft nadE - Ft nadV*, ColE1 *ori*, Amp<sup>R</sup> | [13] |
| pWB301 | *P<sub>LlacOI</sub>::Ec yqhD*, ColE1 *ori*, Amp<sup>R</sup> | This study |
| pWB302 | *P<sub>LlacOI</sub>::Se pnuC\* KA*, ColE1 *ori*, Amp<sup>R</sup> | This study |
| pWB303 | *P<sub>LlacOI</sub>::Ft nadE*, ColE1 *ori*, Amp<sup>R</sup> | This study |
| pWB304 | *P<sub>LlacOI</sub>::Ec pnuC - Se nadR*, ColE1 *ori*, Amp<sup>R</sup> | This study |
| pWB305 | *P<sub>LlacOI</sub>::Ec pnuC - Sc NRK1*, ColE1 *ori*, Amp<sup>R</sup> | This study |
| pDB102 | *P<sub>LlacOI</sub>::Ft nadE - Rs nadV*, ColE1 *ori*, Amp<sup>R</sup> | This study |
| pDB103 | *P<sub>LlacOI</sub>::Ft nadE - Ss nadV*, ColE1 *ori*, Amp<sup>R</sup> | This study |
| pDB104 | *P<sub>LlacOI</sub>::Ft nadE - SynE nadV*, ColE1 *ori*, Amp<sup>R</sup> | This study |
| pLZ301 | *P<sub>LlacOI</sub>::Ft nadE - Hd nadV*, ColE1 *ori*, Amp<sup>R</sup> | This study |

Abbreviations indicate source of genes: *Ec*, *Escherichia coli*; *Se*, *Salmonella enterica*; *Ft*, *Francisella tularensis*; *Sc*, *Saccharomyces cerevisiae*; *Rs*, *Ralstonia solanacearum*; *Ss*, *Synechocystis* sp. PCC 6803; *SynE*, *Synechococcus elongatus* PCC 7942; *Hd*, *Haemophilus ducreyi*. *Se pnuC\* KA* contains mutations compared to the wild type sequence (see Method and supplementary information).

The *yqhD* gene was isolated from *E. coli* BL21 chromosomal DNA by PCR. The resulting PCR fragment was gel purified and assembled into a ColE1 *ori*, AmpR vector backbone by Gibson isothermal DNA assembly method. We used the *yqhD*-harboring plasmid (pWB301) as a control vector in the growth rescue experiments, because it expresses a similar sized-protein to the NMN^+^-producing enzymes using the same promoter, and hence may cause similar growth burden. The gene product of *yqhD* has unrelated function to NMN^+^ biosynthesis.

*E. coli pnuC*, *S. enterica nadR*, *S. cerevisiae NRK1*, *Synechocystis sp nadV*, and *S. elongatus nadV* were isolated by PCR from their respective chromosomal DNA. *F. tularensis nadE\**, *R. solanacearum nadV*, and *H. ducreyi nadV* genes were amplified from *E. coli* codon optimized synthesized DNA templates and assembled as described above.

*S. enterica PnuC\** is generated by site-directed mutagenesis based on the wild type *S. enterica pnuC* gene [29]. The *S. enterica pnuC* gene was isolated by PCR from chromosomal DNA and assembled as discussed above. To perform the KA insertion, which has been shown to enable NMN^+^ transport [29], a sequence of AAAGCA was inserted directly after the 321 base pair, as shown in red text in the Supplemental Information. This resulted in the insertion of a lysine and an alanine residue at the 108 and 109 residue positions. The insertion was introduced by PCR. The subsequent mutant PCR fragments were assembled at discussed above. To generate multi-gene plasmids, the genes were inserted sequentially with a ribosome binding site preceding each gene in the synthetic operon.

Gene sequences are listed in the Supplemental Information.

### Growth-based Screening Platform

Plasmids (selected from pWB301-305) were transformed into *E. coli* strains BW25113 and 72c [27] using the Mix & Go *E. coli* Transformation Kit (Zymo Research).

Overnight cultures were grown in LB medium supplemented with 2 g/L D-glucose, 0.1 mM IPTG, appropriate antibiotics in test tubes at 30 °C while shaking at 250 r. p.m. for 16 hours. For the growth assay, cells were cultured in 1 mL of LB medium supplemented with 2 g/L D-glucose, 0.1 mM isopropyl-β-D-thiogalactopyranoside (IPTG), appropriate antibiotics, and 200 μM of feeding compound, if applicable, in a 2 mL deep-well plate, with square wells, sealed with air permeable film. The medium was inoculated with 1% v/v overnight cultures. Cultures were grown at 42 °C while shaking at 250 r.p.m. Cell growth was monitored by measuring optical density at 600 nm. When applicable, media contained 100 mg/L ampicillin to maintain the plasmids.

### Intracellular NMN^+^ Generation and Quantification

NMN^+^ generation and quantification were performed as previously reported [13]. Briefly, plasmids (selected from pWB203, pDB102-104, and pLZ301) were transformed into *E. coli* strains BW25113 and JW2670-1 (Δ*pncC)* as described previously. Overnight cultures were grown in 2xYT medium containing 0.1 mM IPTG, 2 g/L D-glucose, and appropriate antibiotics for 12 hours at 30°C at 250 rotations per minute (r.p.m.). To cultivate cells for nucleotide analysis, cells were grown in a 50 mL conical tube containing 10 mL of 2xYT media supplemented with 0.5 mM IPTG, 1 mM nicotinamide, and appropriate antibiotics. Cultures were inoculated with 1% v/v overnight culture. Tubes were incubated at 30 °C with shaking at 250 r.p.m. for 4 hours. All media contained 100 mg/L ampicillin to maintain the plasmids.

Cells were processed as previously reported [13]. Briefly, 1 mL of cell culture was pelleted, washed with 1 mL of deionized water, and lysed by resuspension in 1 mL of 95 °C 1% formic acid containing 1 μM of 1-methylnicotinamide as an internal standard. Lysates were quenched in an ice bath before pelleting cell debris. Supernatants were run on a Waters ACQUITY Ultra Performance Liquid Chromatograph with a Waters ACQUITY UPLC CSH C18 column (1.7 μM × 2.1 mm × 50mm). Mobile phases used for separation were (A) water with 2% acetonitrile and 0.2% acetic acid and (B) acetonitrile with 0.2% acetic acid. MS/MS detection was performed by a Waters Micromass Quattro Premier XE Mass Spectrometer. The detailed UPLC and MS parameters were reported previously [13].

### Cell growth with high concentrations of NMN^+^

A plasmid containing the NMN^+^ transporter *S. enterica* PnuC* (pWB302) was transformed into *E. coli* strains JW2670-1 (Δ*pncC*) and MX101 (Δ*pncC* Δ*nadR*). Overnight cultures were grown in 2xYT medium supplemented with 1 g/L D-glucose, 0.1 mM IPTG, and 100 mg/L ampicillin at 30 °C while shaking at 250 r.p.m. for ~14 hours. 1 mL of overnight culture was pelleted in a 1.5 mL microcentrifuge tube and washed twice with 1 mL of 1x M9 salts. The washed cells were resuspended in 1 mL of 1x M9 salts and used for inoculations. For the growth assay, cells were cultured in 1 mL of M9 minimal medium (1x M9 salts, 1 g/L glucose, 1 mM MgSO_4_, 0.1 mM CaCl_2_, 40 mg/L FeSO_4_, 1x A5 trace metals with cobalt) containing 0.5 mM IPTG in a 2 mL deep-well plate, with square wells, sealed with air permeable film. The medium was inoculated with 1% v/v of washed overnight cultures. Cultures were grown at 30 °C while shaking at 250 r.p.m.. Cell growth was monitored by measuring optical density at 600 nm. Media contained 100 mg/L ampicillin to maintain the plasmid.

## Abbreviations

NAD^+^: nicotinamide adenine dinucleotide
NADP^+^: nicotinamide adenine dinucleotide phosphate
P2NA^+^: 3-carbomoyl-1-phenethylpyridin-1-ium chloride
NMN^+^: nicotinamide mononucleotide
NadV: nicotinamide phosphoribosyltransferases
NadE*: nicotinamide mononucleotide synthase
PncC: nicotinamide mononucleotide
NaMN^+^: nicotinic acid mononucleotide
NR: nicotinamide riboside
PnuC: nicotinamide riboside transporter
PnuC*: mutant nicotinamide riboside transporter
Nrk1: nicotinamide riboside kinase from *Saccharomyces cerevisiae*
NadR: nicotinamide riboside kinase (*Salmonella enterica*)
NA: nicotinamide
LC-MS: liquid chromatography-mass spectrometry
NaAD: nicotinic acid adenine dinucleotide
PCR: polymerase chain reaction
IPTG: isopropyl-β-D-thiogalactopyranoside
r.p.m.: rotations per minute

## Declarations

### Ethics approval and consent to participate

Not applicable

### Consent for publication

Not applicable

### Availability of Data and Materials

The datasets used and/or analyzed during this study are available from the corresponding author on reasonable request.

### Funding

H.L. acknowledges support from University of California, Irvine, the National Science Foundation (NSF) (award no. 1847705), and the National Institutes of Health (NIH) (award no. DP2 GM137427). W.B.B. acknowledges support from Graduate Assistance in Areas of National Need fellowship funded by the U.S. Department of Education. D.A. acknowledges support from the Federal Work Study Program funded by the U.S. Department of Education. The content is solely the responsibility of the authors and does not necessarily represent the official views of the National Institutes of Health or the NSF.

### Author Contributions

H.L conceived the research. W.B.B., L.Z., E.K. designed and conducted growth-based screening. W.B.B., D.B., D.A. performed study of prospective NadV homologs. W.B.B. designed and W.B.B., D.A., D.B., L.Z., E.K., performed the intracellular NMN^+^ and NAD^+^ level analysis. W.B.B. and D.A. investigated physiological response to intracellular NMN^+^. All authors analyzed the data and interpreted results. H.L., W.B.B., D.A. wrote the manuscript.

## Acknowledgements

We thank the University of California, Irvine Mass Spectrometry Facility and Dr. Felix Grun for help with LC-MS.

## Conflict of Interest

The authors declare no competing financial interests.

## Corresponding Authors

Correspondence to Han Li.

## Notes

### Competing Interest Statement

The authors have declared no competing interest.

