## Supplementary information for "Metabolic engineering of *Escherichia coli* for optimized biosynthesis of nicotinamide mononucleotide, a noncanonical redox cofactor"

**Supplementary Figure 1:** *Francisella tularensis* NadE*-based Growth Restoration is Not Nicotinamide Feeding Dependent.

**Supplementary Figure 2:** Intracellular $NAD^+$ Decreases in $NMN^+$ Accumulating Strains.

**DNA sequences of genes used in this study**

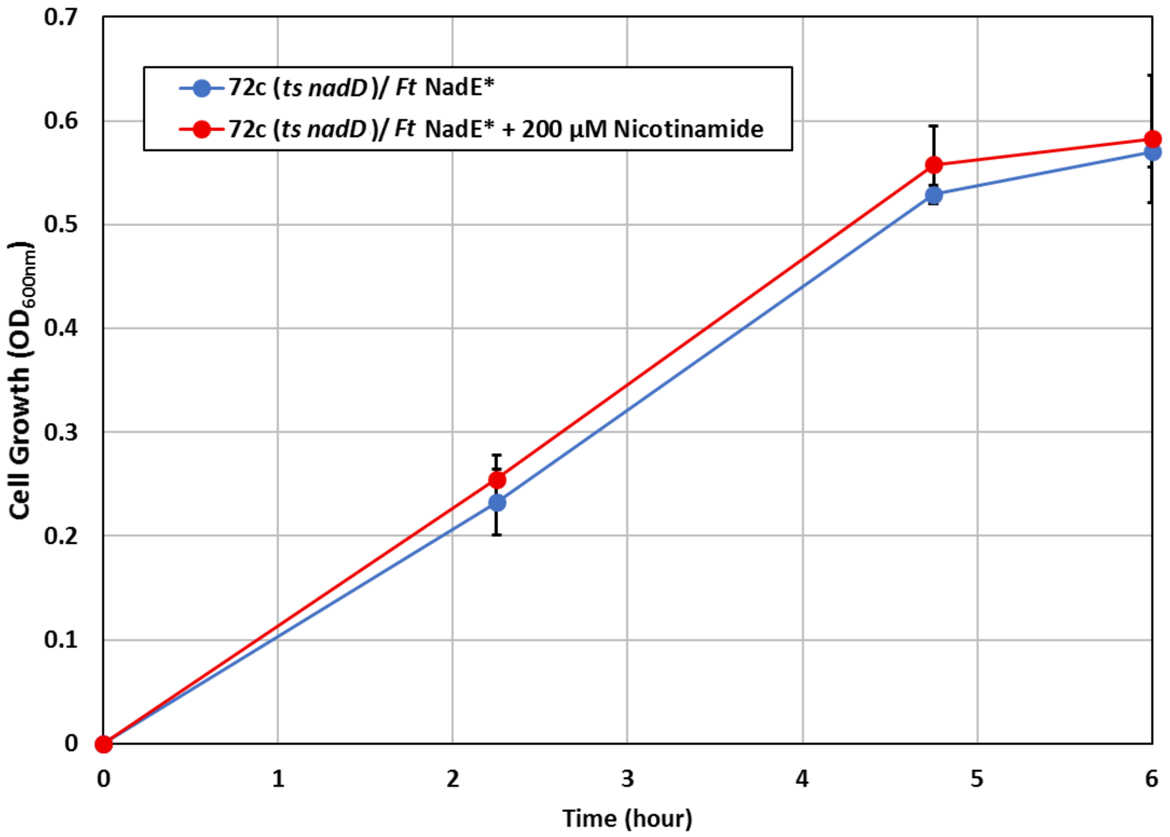


**Supplementary Figure 1: *Francisella tularensis* NadE*-based Growth Restoration is Not Nicotinamide Feeding Dependent.**

A growth restoration platform was used to screen pathways for the efficient generation of nicotinamide mononucleotide ($NMN^+$). The *Escherichia coli* strain 72c [1] contains a temperature sensitive allele of *nadD* (*ts nadD*). As a result, this strain cannot grow at 42 ℃. By overexpressing *Francisella tularensis nadE**, cells are able to produce $NMN^+$, which can then be converted to $NAD^+$, and thus restoring growth. We observed no dependence of growth restoration with feeding 200 µM nicotinamide (NA). This indicates either efficient $NMN^+$ generation can be achieved through channeling the intermediate nicotinic acid mononucleotide ($NaMN^+$) from *E. coli* 's native *de novo* $NAD^+$ biosynthesis pathway, or LB medium used in this experiment already contains sufficient precursors for this pathway. Screening was performed in a deep-well 96-well plate containing 1 mL of LB medium supplemented with 2 g/L D-glucose and 200 µM of NA if applicable. Detailed conditions are described in the Methods section.

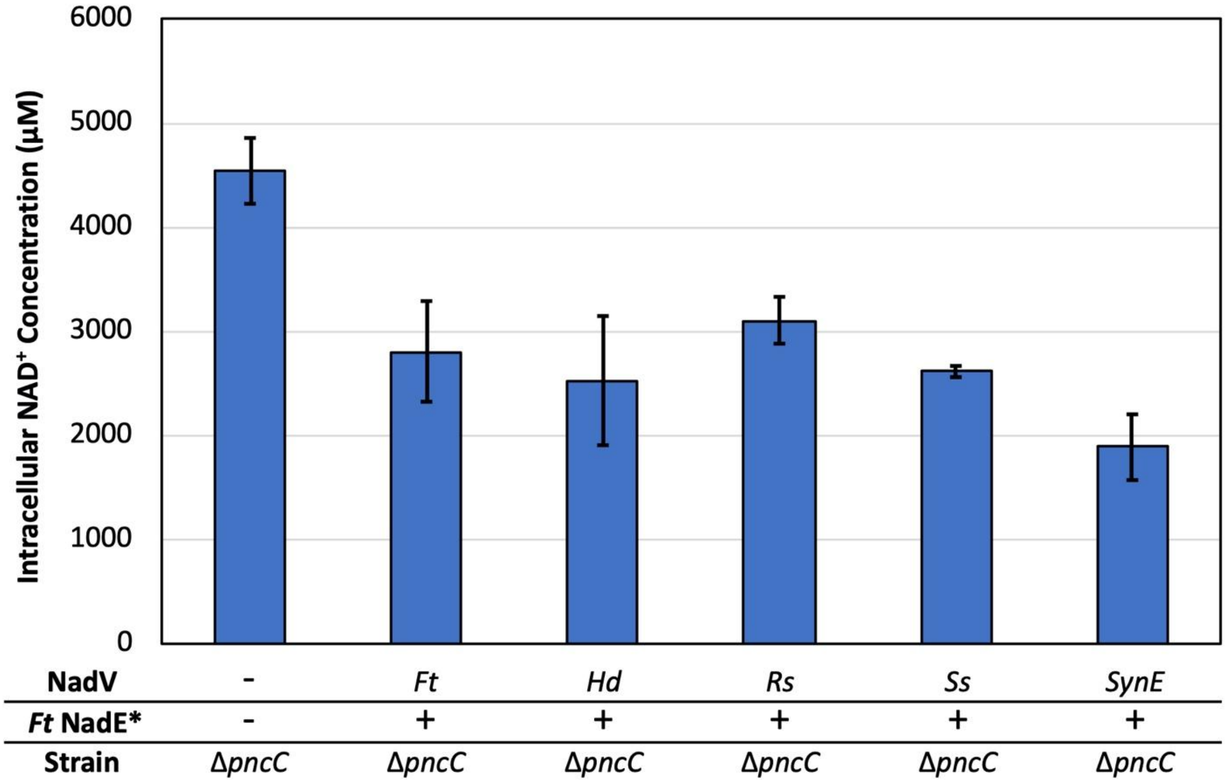


**Supplementary Figure 2: Intracellular NAD$^+$ Decreases in NMN$^+$ Accumulating Strains**
From Figure 3, co-overexpression of NMN$^+$ generating *Francisella tularensis* NadE* and NadVs increases intracellular NMN$^+$ when the NMN$^+$ degrading PncC is disrupted. However, as shown here, NAD$^+$ levels decreased in cells expressing *F. tularensis* NadE* and NadV compared to cells without overexpression. This potentially indicates NMN$^+$ plays a regulatory role in NAD$^+$ biosynthesis. Cells were cultured in 2xYT medium supplemented with 1 mM nicotinamide at 30 ℃ for four hours. Intracellular NAD$^+$ concentrations were determined by UPLC-MS/MS. Detailed conditions and analytical techniques are described in the Methods section.

### DNA sequences of genes used in this study

#### *Escherichia coli* BL21 *yqhD*

ATGAACAACTTTAATCTGCACACCCCAACCCGCATTCTGTTTGGTAAAGGCGCAATC
GCTGGTTTACGCGAACAAATTCCTCACGATGCTCGCGTATTGATTACCTACGGCGGC
GGCAGCGTGAAAAAAACCGGCGTTCTCGATCAAGTTCTGGATGCCCTGAAAGGCAT
GGACGTGCTGGAATTTGGCGGTATTGAGCCAAACCCGGCTTATGAAACGCTGATGA
ACGCCGTGAAACTGGTTCGCGAACAGAAAGTGACTTTCCTGCTGGCGGTTGGCGGC
GGTTCTGTACTGGACGGCACCAAATTTATCGCCGCAGCGGCTAACTATCCGGAAAAT
ATCGATCCGTGGCACATTCTGCAAACGGGCGGTAAAGAGATTAAAAGCGCCATCCC
GATGGGCTGTGTGCTGACGCTGCCAGCAACCGGTTCAGAATCCAACGCAGGCGCGG
TGATCTCCCGTAAAACCACAGGCGACAAGCAGGCGTTCCATTCTGCCCATGTTCAGC
CGGTATTTGCCGTGCTCGATCCGGTTTATACCTACACCCTGCCGCCGCGTCAGGTGG
CTAACGGCGTAGTGGACGCCTTTGTACACACCGTGGAACAGTATGTTACCAAACCGG
TTGATGCCAAAATTCAGGACCGTTTCGCAGAAGGCATTTTGCTGACGCTAATCGAAG
ATGGTCCGAAAGCCCTGAAAGAGCCAGAAAACTACGATGTGCGCGCCAACGTCATG
TGGGCGGCGACTCAGGCGCTGAACGGTTTGATTGGCGCTGGCGTACCGCAGGACTG
GGCAACGCATATGCTGGGCCACGAACTGACTGCGATGCACGGTCTGGATCACGCGC
AAACACTGGCTATCGTCCTGCCTGCACTGTGGAATGAAAAACGCGATACCAAGCGC
GCTAAGCTGCTGCAATATGCTGAACGCGTCTGGAACATCACTGAAGGTTCCGATGAT
GAGCGTATTGACGCCGCGATTGCCGCAACCCGCAATTTCTTTGAGCAATTAGGCGTG
CCGACCCACCTCTCCGACTACGGTCTGGACGGCAGCTCCATCCCGGCTTTGCTGAAA
AAACTGGAAGAGCACGGCATGACCCAACTGGGCGAAAATCATGACATTACGTTGGA
TGTCAGCCGCCGTATATACGAAGCCGCCCGCTAA

#### *Francisella tularensis nadE** (Codon optimized for *E. coli*)

ATGAAAATCGTTAAGGATTTTAGCCCGAAAGAATACTCCCAAAAGCTGGTAAATTG
GTTGAGCGACTCATGTATGAACTACCCGGCCGAGGGATTCGTTATCGGCTTGAGTGG
TGGTATTGACTCAGCCGTCGCGGCCTCATTGGCTGTCAAAACGGGCCTTCCTACGAC
TGCCTTAATTTTGCCGTCCGATAATAATCAGCATCAAGATATGCAGGATGCACTGGA
GTTGATCGAAATGCTTAACATTGAGCACTATACTATCTCGATCCAACCGGCGTATGA
GGCCTTCCTGGCTTCTACACAGAGTTTCACCAATCTTCAGAATAATCGTCAACTTGTC
ATTAAAGGCAACGCCCAGGCTCGTCTGCGCATGATGTATCTGTATGCATACGCCCAA
CAATACAATCGTATCGTCATTGGCACCGACAATGCGTGCGAATGGTACATGGGTTAT
TTCACGAAGTTTGGCGATGGTGCTGCCGACATTCTGCCACTGGTAAACCTTAAAAAG
TCACAAGTTTTTGAGCTGGGTAAATATCTGGACGTTCCCAAAAATATCTTAGACAAG
GCTCCGTCGGCTGGATTGTGGCAAGGGCAAACCGACGAGGATGAAATGGGGGTTAC
CTATCAAGAGATTGACGACTTCTTAGATGGGAAACAGGTTAGTGCCAAGGCCCTGG
AGCGTATCAATTTCTGGCATAACCGCTCGCATCATAAACGTAAATTAGCTTTGACCC
CAAACTTTtaa

***Salmonella enterica pnuC* KA***, red text indicates the KA insertion made to the DNA sequence

ATGGATTTTTTTAGTACGCACAACATACTGATTCATATTCCGATTGGCGCTGGCGGG
TACGATCTCTCGTGGATCGAAGCGGTAGGAACCATCGCCGGCCTGCTCTGTATTTGG
CTTGCCAGTCTGGAGAAGATCAGCAACTACTTTTTTGGACTGGTTAACGTTACCCTG
TTTGCGATTATTTTCTTTCAGATCCAGCTTTATGCCAGCCTGTTGCTGCAACTCTTTTT
CTTTGCCGCCAATATTTATGGCTGGTATGCGTGGTCGCGGCAAACAAAGGATAATCA
AGCCGAGCTTAAAATCCGCTGGCTGCCGTTGCCA**AAAGCA**AAAGCAATGGCATGGC
TGGCGATATGTGTGATAGCTATCGGTTTGATGACGCGATATATCGATCCCGTATTCG
CCGTCCTGACGCGCGTGGCCGTCGCCATTATGCAGATGCTGGGGTTACAGGTGACAA
TGCCCGTACTGCAACCGGACGCTTTCCCGTTCTGGGACTCTTGCATGATGGTGCTGTC
TATCGTGGCGATGATTCTGATGACACGCAAATATGTCGAAAACTGGCTCCTGTGGGT
GATAATCAACGTGATCAGTGTGGTGATTTTTGCTTTGCAGGGCGTCTATGCGATGTC
GCTGGAATATCTGATCCTGACATTTATCGCCGTGAACGGTAGCCGCCTGTGGATAAA
CAGCGCGCGGGAGCGAGGATCGCGCGCTTTCCCGTTAA

***Escherichia coli* BL21 *pnuC***

ATGGATTTTTTTAGTGTGCAGAATATCCTGGTACATATACCAATAGGGGCAGGCGGT
TATGATCTCTCATGGATCGAAGCGGTAGGCACGATCGCCGGGTTGCTGTGTATTGGC
CTTGCCAGTCTGGAGAAGATCAGCAACTACTTCTTTGGCCTGATCAACGTCACCTTG
TTTGGCATTATTTTCTTTCAGATTCAGCTGTATGCCAGCCTGCTATTACAGGTGTTTTT
CTTTGCCGCGAATATTTACGGTTGGTATGCGTGGTCGCGACAAACCAGTCAGAACGA
GGCGGAGTTGAAAATTCGCTGGTTGCCATTGCCGAAGGCACTCAGCTGGTTGGCGGT
TTGCGTTGTTTCGATTGGTCTGATGACGGTATTTATCAATCCGGTGTTTGCATTTTTG
ACCCGCGTGGCAGTCATGATCATGCAAGCATTAGGATTACAGGTTGTGATGCCTGAA
CTGCAACCGGACGCTTTCCCGTTCTGGGATTCATGCATGATGGTGTTATCTATCGTGG
CAATGATTCTGATGACGCGTAAGTATGTGGAAAACTGGCTGTTGTGGGTGATTATTA
ACGTGATTAGCGTCGTTATTTTTGCACTTCAGGGCGTTTACGCCATGTCTCTGGAGTA
CATCATCCTGACCTTTATTGCGCTCAACGGCAGCCGGATGTGGATCAACAGCGCACG
TGAAAGAGGCTCACGCGCGCTGTCCCATTAA

***Salmonella enterica nadR***

ATGCGATTTTTCCAGCAGGAGGCTCTTGTGTCATCGTTCGACTATCTCAAAACCGCG
ATTAAGCAGCAAGGTTGCACTCTGCAACAGGTGGCTGACGCCAGCGGTATGACCAA
GGGATATCTGAGTCAGTTACTTAACGCCAAAATCAAAAGCCCCAGCGCGCAAAAAC
TGGAGGCGCTACACCGTTTTCTCGGGCTGGAGTTTCCCCGCCGACAGAAAAACATTG
GCGTGGTGTTCGGTAAATTTTATCCATTGCATACCGGACACATCTACTTGATCCAGC
GCGCCTGTAGCCAGGTGGATGAGTTGCACATCATTATGGGATATGACGATACGCGC
GACCGCGGGCTGTTTGAGGATAGCGCCATGTCGCAGCAGCCCACCGTGTCGGATCG
CCTGCGCTGGTTATTGCAAACCTTCAAATACCAAAAAAATATTCGCATCCACGCCTT

TAATGAAGAGGGGATGGAGCCTTATCCGCATGGCTGGGACGTCTGGAGCAACGGCA
TTAAAGCGTTTATGGCAGAGAAGGGAATACAGCCGAGCTGGATCTACACTTCCGAA
GAGGCTGATGCGCCGCAGTATCTTGAGCATTTAGGGATTGAGACGGTGCTGGTCGAT
CCTGAACGCACGTTTATGAATATCAGTGGGGCGCAAATCCGCGAAAATCCGTTTCGT
TACTGGGAATATATTCCTACCGAAGTGAAGCCGTTTTTCGTGCGTACCGTCGCGATT
CTGGGCGGGGAATCAAGCGGCAAGTCTACGCTGGTCAATAAGCTCGCCAATATTTTT
AATACCACCAGCGCCTGGGAATATGGCCGCGACTATGTCTTTTCGCATCTGGGCGGC
GATGAGATGGCGTTACAGTATTCCGACTACGATAAAATTGCGCTGGGCCATGCGCA
ATATATTGATTTCGCAGTGAAATATGCGAATAAAGTGGCGTTTATCGATACCGATTT
CGTCACCACCCAGGCATTTTGCAAAAAATACGAAGGACGCGAGCATCCCTTTGTCCA
GGCGCTGATCGACGAGTATCGCTTCGACCTGGTGATTTTGCTGGAGAATAATACGCC
GTGGGTAGCTGACGGACTGCGAAGCCTGGGCAGTTCAGTGGATCGCAAAGCGTTCC
AGAACCTGCTGGTCGAGATGCTGAAAGAGAACAACATTGAGTTCGTTCACGTTAAA
GAGGCTGATTACGATGGTCGCTTTTTGCGCTGTGTGGAACTGGTGAAAGAGATGATG
GGCGAGCAGGGATAA

***Saccharomyces cerevisiae* BY4741 *NRK1***

ATGACTTCGAAAAAAGTGATATTAGTTGCATTGAGTGGATGCTCCTCCAGTGGTAAG
ACGACAATTGCGAAACTTACAGCAAGTTTATTCACGAAGGCTACATTAATTCATGAA
GATGACTTTTACAAACATGATAATGAAGTGCCAGTAGATGCTAAATATAACATTCAA
AATTGGGATTCGCCAGAAGCTCTTGATTTTAAACTTTTCGGTAAAGAATTAGATGTG
ATCAAACAAACTGGTAAAATAGCCACCAAACTTATACACAATAACAACGTAGATGA
TCCCTTTACAAAGTTCCACATTGATAGACAAGTTTGGGACGAGTTAAAGGCTAAGTA
TGACTCTATTAATGACGACAAATATGAAGTTGTAATTGTAGATGGGTTTATGATTTT
CAATAATACTGGAATATCAAAAAAATTTGATTTGAAGATATTAGTGCGTGCTCCCTA
TGAAGTACTAAAAAAAAGGAGGGCTTCCAGAAAAGGATACCAGACTTTGGATTCTT
TCTGGGTGGATCCGCCGTATTATTTCGACGAATTTGTGTATGAATCTTATCGTGCAAA
TCATGCGCAGTTATTTGTTAATGGAGACGTAGAAGGTTTACTAGACCCAAGGAAGTC
AAAGAATATAAAAGAGTTCATAAATGATGATGACACTCCAATTGCGAAACCTTTAA
GCTGGGTGTGCCAAGAGATTCTAAAGCTTTGTAAGGATTAG

***Francisella tularensis nadV* (Codon optimized for *E. coli*)**

ATGTCGTTTGATAACCTGCTGTTGATGACGGATTCCTATAAACATAGTCACCGTTAC
CAATATCCTCGCGATACCCATTATCTGCATTTTTATCTGGAATCACGCGGGACCGCT
AACAAGGATCTGGGCAACTATACGAAATTCTTCGGATTGCAGTATTACGTTAAAAAG
TATCTTTCCCAACCCATTACCCAGCAGATGATCGACGATGCAGAGAAGATCTTACTT
GCCCACGGGCTTCCGTTCTACCGTAGTGGGTTCGAGAAGATCCTTAATAATTATAAC
GGATACCTGCCGATTCGTATCCGTGCCGTGCGTGAAGGTAGTTTAATCCCGCTGCAT
AATGTATTAATGACGATTGAGTCGACGGACGAAGAGCTTTTCTGGCTTCCGGGCTTC
GTAGAAACTCTGCTGTTGAAGGTATGGTACCCAACGACTGTAGCTACGATTAGCTTT
AATATCAAACAACTGATTAAACGTTACTTGTTGGAGACGGCAGACTCGCTTGATAAG

TTAGACTTTATGTTGCATGACTTTGGATACCGCGGTGTCTCTAGTGAGGAGTCAGCA
GGTATTGGGGGGGCCGCGCATCTGACCAATTTTTTGGGCACCGATACATTAGCGGCC
CTTCATGTTTGTAAAGAGTTCTATGCGGAGGACATGGCAGGATTTTCCATCCCTGCG
TCGGAACATTCAACTATGACTAGCTGGGGCGTGGGGACCGAGTGTGAGCGCGAAGC
GTTTGAAAATATGATTGCGCAGTTCGGTGACTCTTCGGTCTTATATGCTTGTGTCTCT
GACTCATGGGACTTTAAAAAAGCGATCCAGACCTGGGTAGACTTGAAAGACCGCGT
TACCGCCAAAAAGGCGAACTTAGTAATCCGTCCAGACAGTGGCGACGCCGTAGATA
ACATTTTGTACGCGCTTTATGAACTTGACAAAGGGTATGGATCACGTTTAAATAGTA
AGGGGTACAAAGTTTTAAACAATGTAGCACTTATTCAAGGGGACTCTGTTTCTATTT
CGTTAGCGAAGAAAGTTTTAGAGGCCATGAAAATTCAAGGCTACTCCGCAGAGAAC
ATTGCATTCGGGATGGGAGGGGCTCTTCTGCAAGGGAACTACGAATCGTCGATCAA
CCGCGACAGCTTCAAATTCGCAATCAAATGTTCTGCTATTATGCGCGGTAATACTTT
AATCGGCGTTAAGAAGGAGCCAATTACCGATCTTGCTAAGAAATCAAAACAGGGTC
GTTTGGATCTTATTAAGGACGCGAAAGGAAATTACAAAACGATCGTACTGGACGAC
TCGTATGCGTTAGGTGAGTATCATCCGGAATCTCAATTGCAAACCTACTATGATAAT
GGCGAGATCAAGTTTGAACAGAGCCTTGCCCAGATCCGTAATTACACAAATTAA

***Ralstonia solanacearum nadV* (Codon optimized for *E. coli*)**

ATGCAGAACGACCTGCCTGGTTTGTCCGCTATCCTTAGCAACCCAATCTTAAATACC
GACAGTTACAAGGCGTCGCATTACCTGCAATACCCAGCCGGTACTTCGGCGATGTTC
TCCTACGTAGAATCCCGTGGAGGTCGTTATGATCGTACCGTTTTCTTCGGACTTCAAA
TGCTGGCAAAGGAATACTTATGCCGTCCTATTACCCCTGCTATGATCGATGCTGCCC
GCGGGTTTTTCGCAGCACACGGGGAGCCGTTTAACGAGGCGGGATGGCGTTATATTG
TTGCCCGTTATGATGGCTATCTGCCCGTACGTATCCGTGCGGTTCCCGAGGGGTCAG
TGGTACCTAATCACAACGTGCTGATGACAGTCGAATGTGACGATCCTGAAGTTTTCT
GGCTTGCGTCATATCTGGAAACTATGTTATTGCGCGTGTGGTATCCGATTACAGTTG
CGACCCAGAGCTGGCATCTGCGTCAACTTGTCCACCGCTACCTGGAGCAAACAAGT
GATGACCCAGGACAGTTGCCATTCAAGGTTCATGATTTCGGTGCTCGCGGTGTATCT
AGCGCGGAAAGTTCGGCTATTGGGGGAGCAGCTCACCTTGTGTCTTTCATGGGTAGT
GACACGGTTTTGGGTGTGGCCGCCGCAAACCTGTATTACAATGCTCAAATGGCCGCG
TTTTCTGTACCCGCGGCGGAGCACAGTACGATTACAGCCTGGGGACGTGCCGGGGA
AGCAGATGCGTATCGTAATATGTTACGCCAATTCGGTAAACCTGGTGCGATCGTGAG
TGTTGTCAGTGACAGTTATGACTTATTCGCCGCGCTTCGCCTGTGGGGAGGGGAATT
ACGCCAGGCAGTCATCGACTCTGGGGCTACGCTTGTCGTACGTCCCGATTCTGGCGA
CCCTCGCTCCATTGTTCTTCAGACAGTCCGCGCGCTTGATGCTTCATTTGGAGCAACA
GTGAACGGGAAAGGGTACCGTGTCCTGAACCACGTCCGCGTCATTCAAGGCGATGG
AATTAATGCAGCATCGATCGAGGCAATTCTTGCCGAGTTAGAGGCTGCGGGATATGC
GGCGGATAACATTGTATTCGGGATGGGAGGTGCCCTGTTACAACAATTAAACCGCG
ACACACAGCGCTTTGCAATGAAGTGCTCAGCAGTCCGTGTTGACGGGGCGTGGCGT
GAAGTCTGTAAAGACCCGGTGACCGACGCGGGGAAACGTTCTAAGAAAGGACGTCT
TACACTTTTGCGCAACCGTGTGAGCGGGGAGTACGCCACAGCCACTTTGCCCTTGGC

CTGGGATGATCGCCGCATCGAGGGGGAATGGGAGGATGCTCTGGTGACGGTATTCG
AGAATGGGCGTCTTTTACAGGATGTCAGCCTTGACGCGGTCCGCGCGCGCTCAAG
CCCATGAGTTGGCACCCGCCCTTGTCGACTGA

***Synechocystis sp.* PCC 6803 *nadV***

ATGAATACTAATCTCATTCTGGATGTGGACTCCTATAAAGTGAGCCACTGGTTGCAG
TATCCTCCTGACACAACGGCAATGTATTCCTATGTGGAAAGTCGTGGGGGAAGGTAT
CCTGTCACTGTCTTTTTTGGTCTCCAATACATTTTAAAGCGGTATCTGACTCAATCCA
TTGAACCCTGGATGGTGGAGGAAGCTAATCGCCTTTTGACAGCCCATGGCTTACCTT
TCAACTATGGCGGTTGGCGATACATTGCGGAGGATTTGCAGGGTCGTTTACCTGTAC
GTATTAAGGCGGTTCCAGAGGGCTCGGTCATCCCGGTTCATAATGTTTTGATGACAG
TGGAATCCACGGACCCAAAGGTTTTTTGGTTAGTTTCCTGGTTAGAAACTTTGTTGAT
GCGGGTTTGGTATCCCATTACGGTGGCAACCCAGAGTTGGCATTTAAAACAACGCAT
CTATCAATCCCTATGCCGTACTGCGGATGATCCTGATGGTGAAATCAATTTTAAACT
CCACGATTTTGGGGCCCGGGGGGTTTCTAGTGGTGAATCGTCCGGCATTGGCGGACT
GGCTCACTTAGTTAATTTCCAAGGTTCTGACACAGTAAAGGCCCTGGTGTATGGGCA
GCAATATTACAACTGCCCCATGGCGGCCTATTCGATTCCCGCCGCAGAACATTCCAC
CATTACAGCTTGGGGAAGGGAAGGGGAAGTTTTGGCCTATGAAAATATGTTGACCC
AGTTTGCCAAGCCAGGGTCGGTGTTGGCGGTGGTTTCCGATTCCTATGATCTCTGGA
ATGCCATTGACCATCTCTGGGGCGATCACCTAAGGGCACAGGTGCTTGATTCGGGGG
CTACGGTGGTTATCCGTCCGGATTCAGGTGACCCGGTGGCCATTGTGGCCCAAACTT
TGGAACGGTTGGAGGCTTGTTTTGGCAGCACCCTCAACAGTAAGGGCTTTCGAGTTC
TAAATGCTGTGCGGGTTATCCAAGGGGATGGGGTTGATGAAGAGAGTATCAGCGCC
ATTCTAGAGAAGACTGAGAGCCTTGGCTTTAGTACTACTAATTTAGCTTTTGGTATG
GGGGGAGCTTTGTTGCAAAAGGTGAATCGGGATACCCAAAAATTTGCCATGAAGTG
CAGTGAGGTAACGGTGGAGGACAAGGCGATCCCTGTTTATAAAGACCCTGTTACTG
ATCCTGGTAAAACTAGCAAAAAGGGGCGATTATCCCTGGTTAAAACTGACTCTGGTT
ATGGCACTGTACCCACTTCTTCTGAGGATTTATTGCAGGTTGTCTATGAAAATGGAC
ATTTACTGCAAGACCAATGCTTGGATGCTATTCGTCAACGAGCCTGGCCATTAATCA
GGGTCAATGTTCCCGCAAGCTAG

***Synechococcus elongatus* PCC 7942 *nadV***

ATGGACCTCAATCTTCTGTTCGATACCGACTCATACAAAGTCAGCCACTGGCTGCAA
TATCCTGCCGATACGACTGCGATCGGAGCTTATTTAGAAAGCCGGGGTGGAGATTGC
TCGCACACGCTCTTTTTTGGCTTGCAATATCTACTACTGCGTTATTTCTTCCAGCCAA
TCACTAGCGCTGACATTCAAGAAGCCGCCGCGCTGTTTCAAGCGCATGGGCTGCCTT
TCAATCAAGCGGGCTGGCAACGAGTTTGCGATCGCTATGGCGGGTATTTACCTTTAC
GAATTCGGGCTGTCCCGGAAGGTAGCCTTGTCCCCACCGGCAATATCTTGCTGACAG
TGGAATCGACCGATCCTGAATTGGCTTGGCTGGCCACTTGGGTTGAGACACTACTGC
TGCGGGTTTGGTATCCGATAACTGTGGCTACACGCAGTTGGCAGCTTCGGCAAATCA

TTCAGCAAGCGCTGGAGCAATCAGCCGAAAATCCAGCAGCTGAAATTGACTTCAAA
CTGCATGACTTTGGATCACGCGGGGTATCGAGCCAAGAAAGTGCTGCAATCGGCGG
GCTGGCTCATTTGGTCAACTTTCAAGGCACTGATACGATCGCTGCGTTACTGGCAGG
ACAGCGCTATTACGATTGCGCGATCGCTGGCTTTTCGATTCCGGCGGCGGAGCATTC
AACGATTACGGCTTGGGGCCCATCGGGTGAGTTAGATGCTTACCGCAATATGCTCGA
TCGCTTTGCAAATCCGGGATCTGTGGTGGCTGTTGTATCGGACTCCTATGATCTCTGG
CATGCCGTCGATCAGCTTTGGGGTGAGGATCTCCGCGATCGCATTTTGCAATCGGGA
GCAACCGTTGTCATTCGGCCTGACTCAGGCAATCCTGAGCAGATTGTGCCGGAATTA
CTGCGTCGTTTGGCCGCTAAGTTCGGCTGCGATCGCAATCAGAAGGGTTATCAAGTT
TTGCGATCGGTGCGGGTGATTCAGGGCGATGGGATCACAGTGGACAGTCTGCCCAA
AGTTCTGCAAGCGGTTATGGCCGCTGGCTTTAGTGCCAGTAATGTCGCTTTTGGCAT
GGGTGGCGGGCTGTTGCAGCAGGTCAATCGCGATACCCAACGCTTTGCCTACAAGTG
CAGCTGGATCGAGCGATCGGGACAAGTGATTCCCATTTGCAAGCGACCAGCCACGG
ATCTGCGCAAGGCTAGCAAAGCAGGACGCTTGGATTTAATTCGCGATCGCGAGGGG
CAATACCGAACAGTCTCGTTACTGACGTCAGAGCCTGACCCGCAATCCTGCCTGCAA
ACGGTGTTTGAAAATGGTGCGATCGTGCGGCGACAAAGCTTGCAGGAAATCCGCGA
TCGCGCTCGTTCTGAGACACGCTAG

***Haemophilus ducreyi nadV* (Codon optimized for *E. coli*)**

ATGGATAATTTATTAAATTACTCTTCGCGTGCTTCGGCCATTCCGTCGCTGTTGTGTG
ACTTTTATAAGACATCGCACCGCATTATGTATCCCGAGTGTAGTCAAATCATTTACTC
GACCTTCACACCACGTTCCAATGAACAAGCTCCATACCTGACACAAGTTGTCTCATT
CGGCTTTCAGGCATTTATTATTAAGTACCTTATTCATTACTTCAACGACAATTTTTTC
TCACGCGATAAACATGATGTAGTTACTGAATACTCGGCCTTTATCGAGAAGACTCTT
CAATTAGAAGATACCGGAGAGCATATCGCGAAGCTGCACGAATTGGGCTACCTTCC
GATTCGTATCAAGGCAATCCCCGAGGGGAAAACCGTGGCAATCAAGGTCCCTGTAA
TGACCATCGAGAATACCCATAGTGACTTCTTTTGGTTAACCAATTACTTGGAAACAT
TGATCAACGTTTCGTTGTGGCAGCCGATGACCTCTGCCTCGATTGCATTCGCATATCG
TACTGCCCTTATCAAGTTTGCGAATGAAACGTGCGATAATCAAGAGCATGTCCCCTT
TCAGAGCCACGATTTTTCTATGCGTGGAATGAGTTCCCTGGAATCAGCCGAAACATC
TGGTGCTGGACACTTAACTTCGTTCCTGGGTACGGACACCATTCCGGCGCTTTCCTTT
GTCGAGGCTTATTACGGCAGTAGTAGCCTTATTGGTACATCGATCCCTGCAAGTGAA
CATTCTGTGATGTCCTCACATGGAGTTGACGAGTTAAGTACTTTCCGCTACTTAATGG
CTAAGTTCCCGCATAACATGCTGAGTATCGTGAGTGATACGACTGACTTTTGGCACA
ATATCACGGTCAACCTTCCGTTATTGAAGCAAGAAATCATTGCGCGTCCAGAGAATG
CTCGTCTGGTTATCCGCCCTGATTCAGGGAACTTCTTTGCAATCATCTGCGGAGACCC
AACCGCGGACACCGAACACGAACGCAAGGGTCTTATCGAATGTTTGTGGGACATCT
TCGGAGGGACTGTCAACCAAAAGGGTTATAAAGTTATCAATCCACACATCGGAGCG
ATCTATGGGGACGGCGTAACCTATGAAAAGATGTTCAAGATCTTGGAAGGTCTTCAA
GCCAAAGGATTTGCCTCCAGCAACATTGTATTTGGCGTGGGGGCGCAGACTTACCAG
CGTAATACTCGCGATACATTGGGATTTGCACTGAAGGCCACCTCTATTACTATTAAT

GGAGAGGAGAAGGCAATTTTCAAGAATCCCAAAACAGATGATGGCTTTAAGAAGAG
CCAGAAGGGACGCGTCAAAGTTCTTTCCCGCGATACTTACGTAGATGGCTTAACAAG
CGCTGACGACTTTAGCGACGACCTTCTGGAGCTGCTTTTCGAGGACGGCAAGCTTTT
ACGCCAGACCGATTTCGACGAAATTCGTCAAAATCTGCTTGTGTCCCGTACCACTTT
GTAA
